# Risky Decision-Making Predicts Dopamine Release Dynamics in Nucleus Accumbens Shell

**DOI:** 10.1101/572263

**Authors:** Timothy G. Freels, Daniel B. K. Gabriel, Deranda B. Lester, Nicholas W. Simon

## Abstract

The risky decision-making task (RDT) measures risk-taking in a rat model by assessing preference between a small, safe reward and a large reward with increasing risk of punishment (mild foot shock). It is well-established that dopaminergic drugs modulate risk-taking; however, little is known about how differences in baseline phasic dopamine signaling drive individual differences in risk preference. Here, we used *in vivo* fixed potential amperometry in male Long-Evans rats to test if phasic nucleus accumbens shell (NACs) dopamine dynamics are associated with risk-taking. We observed a positive correlation between medial forebrain bundle-evoked dopamine release in the NACs and risky decision-making, suggesting that risk-taking is associated with elevated dopamine sensitivity. Moreover, “risk-taking” subjects were found to demonstrate greater phasic dopamine release than “risk-averse” subjects. Risky decision-making also predicted enhanced sensitivity to nomifensine, a dopamine reuptake inhibitor, quantified as elevated latency for dopamine to clear from the synapse. Importantly, this hyperdopaminergic phenotype was selective for risky decision-making, as delay discounting performance was not predictive of phasic dopamine release or dopamine supply. These data identify phasic NACs dopamine release as a possible therapeutic target for alleviating the excessive risk-taking observed across multiple forms of psychopathology.

**Significance Statement:** Excessive risky decision-making is a hallmark of addiction, promoting ongoing drug seeking despite the risk of social, financial, and physical consequences. However, punishment-driven risk-taking is understudied in preclinical models. Here, we examined the relationship between individual differences in risk-taking and dopamine release properties in the rat nucleus accumbens shell, a brain region associated with motivation and decision-making. We observed that high risk taking predicted elevated phasic dopamine release and sensitivity to the dopamine transporter blocker nomifensine. This hypersensitive dopamine system was not observed in rats with high impulsive choice, another behavior associated with substance use disorder. This provides critical information about neurobiological factors underlying a form of decision-making that promotes vulnerability to substance abuse.

## Introduction

Multiple factors contribute to the transformation of reward value during economic decision-making. For example, some rewards are accompanied by risk of an aversive event, which “discounts” the value of the reward (Simon et al., 2009; Andrade and Petry, 2012). Excessive risky decision-making is prevalent in substance use disorder (SUD), and enhances likelihood of drug relapse (Bechara et al., 2001; Brand et al., 2008; Brevers et al., 2014; Verdejo-Garcia et al., 2018). Therefore, understanding the neurobiological factors that drive individual differences in decision-making may have utility for precise medical treatment for vulnerable individuals.

The Risky Decision-making Task (RDT) models risk-taking in rats by measuring preference for a small, safe reward or a large reward accompanied by the risk of foot shock (Simon et al., 2009). Individual differences in this task predict several cognitive and motivational phenotypes associated with vulnerability to substance use disorders, with risk-preferring rats demonstrating elevated cocaine self-administration, impulsive action, nicotine sensitivity, and sign-tracking (Mitchell et al., 2014; Olshavsky et al., 2014; Gabriel et al., 2019). Therefore, quantifying risk-taking in RDT appears to be an effective method of detecting a cluster of behavioral phenotypes associated with vulnerability to SUD (Goldstein and Volkow, 2002).

Dopamine release in the nucleus accumbens (NAC) is a canonical mechanism involved in the valuation of rewards and reward-predictive stimuli (Mogenson et al., 1980; Floresco et al., 2003; Sombers et al., 2009). Manipulating dopamine transmission alters multiple rodent assessments of risky decision-making, including RDT. Systemic amphetamine administration consistently reduces risky decision-making (Simon et al., 2009, 2011; Mitchell et al., 2011; Orsini, Willis, Gilbert, Bizon, & Setlow, 2016), whereas cocaine reduces sensitivity to changing risk levels. Chronic exposure to dopaminergic drugs, which causes long-lasting changes in dopamine activity (Robinson and Berridge, 1993), shifts decision-making toward greater risky decision-making (Mitchell et al., 2014). Furthermore, risk-taking in RDT predicts dopamine receptor expression in NAC shell (NACs), but not core (Simon et al., 2011). However, little is known about how individual differences in risk-taking are related to functional dopamine release dynamics in NACs.

Fixed potential amperometry, also known as continuous amperometry, is an ideal neurochemical tool for assessing patterns of dopaminergic activity *in vivo*, given its high temporal resolution of 10,000 samples/s (Benoit-Marand et al., 2007). Stimulating projections from medial forebrain bundle (MFB) to NAC mimics biologically relevant phasic dopamine release, a critical component of motivated behavior (Ikemoto & Panksepp, 1999; Saddoris et al., 2015). Here, we characterized rats in RDT, then assessed how individual differences in risk-taking predict evoked NACs dopamine release, supply, half-life, and sensitivity to the dopamine transporter inhibitor nomifensine. In addition, we compared NACs dopamine release dynamics to delay discounting, another form of decision-making associated with SUD vulnerability (Garavan and Hester, 2007; Perry and Carroll, 2008; Bickel et al., 2014).

## Materials and Methods

### Subjects

Male Long-Evans rats aged approximately 90 days old (*n* = 20) were obtained from Envigo. Rats arrived pair-housed and kept on a 12-hour light/dark cycle beginning with lights off at 8am. Rats had ad libitum access to water, and were food restricted to 90% free feeding baseline weight to increase motivation to pursue the reward in behavioral tasks. Pairs were separated if fighting or food domination was observed. All experiments were approved by The University of Memphis Institutional Animal Care and Use Committee.

### Behavioral Apparatus

Behavior and decision-making processes were measured in Med Associates (Fairfax, VA) operant conditioning chambers equipped with one retractable lever on either side of an illuminable food trough, pellet dispenser, and shock grate. Tailor-made Med Associates codes were used to run all behavioral protocols.

### Risky Decision-Making Task

Rats were run in the RDT to determine individual preference for risky rewards (Simon and Setlow, 2012). Critically, the order of RDT and delay discounting (DD) was counterbalanced across subjects, with some rats beginning with RDT, and others beginning with DD. An approximate timeline for experimental procedures is shown in Figure 1 A.

**Figure 1.**
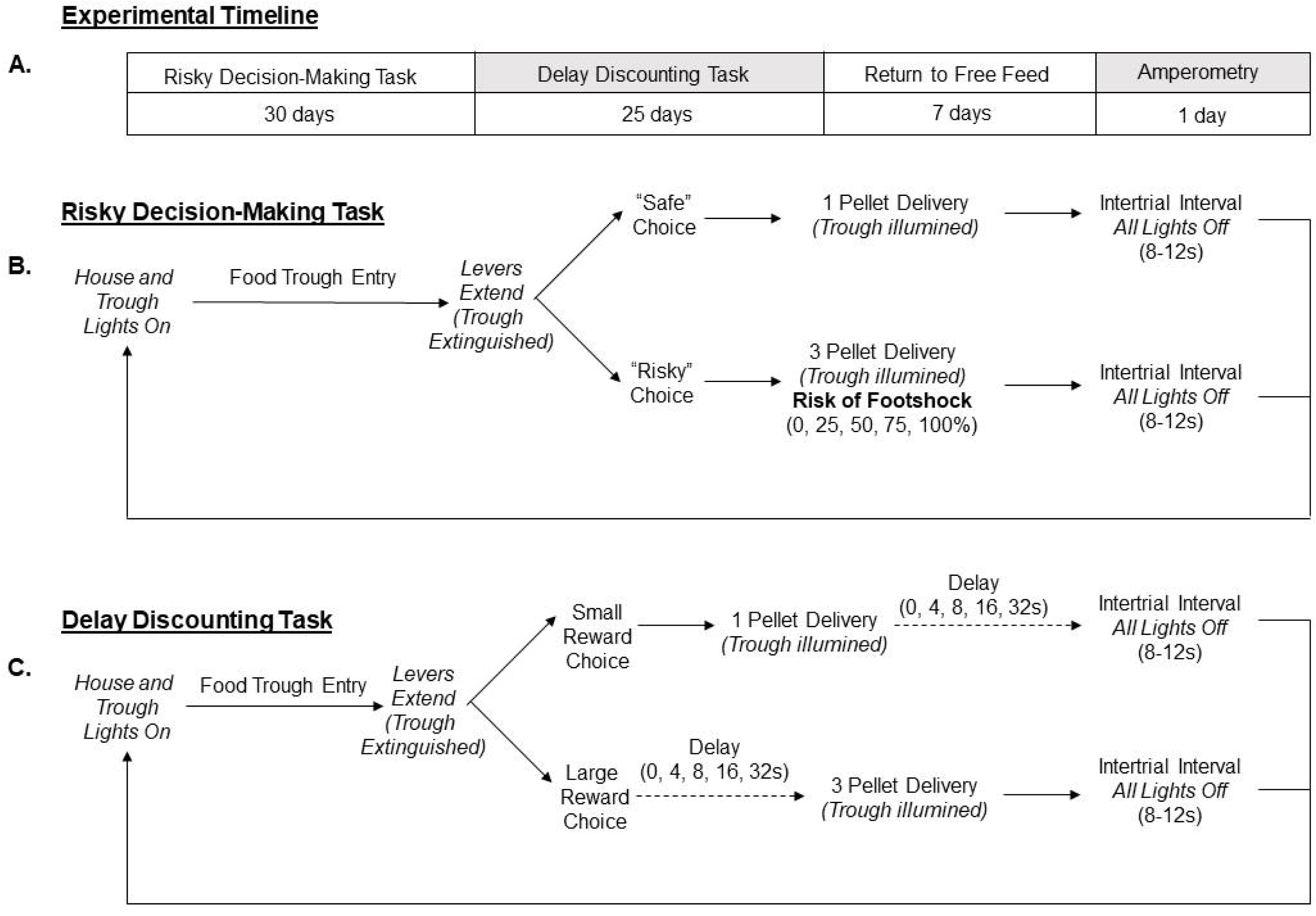
General experimental design. *A.* Experimental timeline for procedures. *B.* Schematic for the risky decision-making task. *C.* Schematic for the delay discounting task.

Initial shaping was adapted from past experiments (Simon et al., 2007). Briefly, rats were trained to associate the food trough with food delivery, initiate lever extension with a food trough entry, and associate both levers with food pellet delivery. Then, rats learned to discriminate between simultaneously presented levers that yielded either large (3 pellet) or small (1 pellet) rewards, with lever identity (right vs left of food trough) counterbalanced across subjects. Upon demonstrating consistent preference for the large reward (>75% choice of it in a session), they began training in the RDT. This consisted of 5 blocks of 18 trials, totaling 90 trials per session. Each block began with 8 forced trials (single lever) followed by 10 choice trials (dual lever). Forced choice trials served to establish new risk contingencies upon initiation of each new block. During forced choice trials, lever presentation was pseudorandomized with the same lever never extended more than twice in a row. After 4 forced choice trials on each lever, both levers were extended upon trial initiation, offering rats a choice between a small safe or large, risky reward. Each trial began with simultaneous illumination of house and trough lights. Rats had 10 seconds to initiate a trial via food trough entry, which extinguished the food trough light and extended either 1 or both levers, depending on trial type.

Selection of the safe lever resulted in the delivery of a single food pellet. A risky lever press delivered 3 food pellets with the risk of a 1 second footshock. The risk of punishment escalated across blocks (0, 25, 50, 75, and 100%). Upon lever press and pellet delivery/shock, levers retracted and the food trough light illuminated. After food was collected or ten seconds passed, the food trough and house lights were extinguished and an intertrial interval (ITI, 10 ± 4s) preceded the next trial. Failure to initiate a trial or press a lever within 10 s of instrumental activation resulted in the trial being marked as an omission and proceeding to the ITI (Figure 1 B).

The initial RDT sessions did not include risk of shock, allowing rats to acquire magnitude discrimination (1 vs. 3 pellets). Once subjects demonstrated a minimum of 75% preference for the large reward within a session, risk of shock was added to this reward. Shock intensity increased from .15 mA to .35 mA across multiple sessions until rats showed a clear preference for the large reward and no complete risk aversion. Using .35 mA for the initial shock amplitude can induce an excessive number of omitted trials and a strong bias away from the large reward during task acquisition (Simon and Setlow, 2012). After reaching .35 mA, rats continued to perform in the RDT until stability was achieved over 5 days, defined as a lack of effect of day or day x block interaction following a repeated measures ANOVA.

### Impulsive Choice

Impulsive choice was assessed using a DD task (modified from Simon et al., 2007). As with RDT, rats chose between a large (3 pellet) and small (1 pellet) reward. However, in the DD task, the large reward was delivered after a delay that increased throughout the session. The session began with a 0 s delay preceding the large reward, which increased to 4, 8, 16, and 32 s over 5 blocks of 10 choice trials each. Each block was preceded with a forced choice trial on each lever to establish delay duration (rats required 20 days of training to acquire DD). The delay before large reward delivery was added to the ITI after each small reward choice to keep trial length consistent between trial types (Figure 1 C). The total percent choice of the delayed reward over a final 5 day average served as a measure of impulsive choice, or willingness to wait for a larger reward.

### Fixed Potential Amperometry

Phasic dopamine release in the NACs was measured using *in vivo* fixed potential amperometry. Rats (*n* = 19) were anaesthetized using two i.p. injections of urethane (totaling 1.5g/kg). They were then placed on a heating pad to maintain a constant body temperature (38 ± 0.1 °C) and fixed into a stereotaxic frame, which held the drill, a 3 electrode recording system, and a stimulating electrode. A concentric bipolar stimulating electrode was placed into the medial forebrain bundle (MFB; coordinates in mm from bregma: AP −4.2, ML +1.8, and DV −7.8 mm from dura; (Paxinos and Watson, 1997). Projections from the ventral tegmental area converge in the MFB in route to striatal regions (Mogenson et al., 1980). Electrical stimulation of the MFB has been shown to evoke dopamine release in the NAC in previous studies (Dugast et al., 1994; Fielding et al., 2013). A combination reference and auxiliary electrode coupling was placed in cortical contact contralateral to the stimulating electrode. Finally, a carbon fiber recording microelectrode (active recording surface of 500 µm length × 7 µm o.d.) was placed in an area estimated as the NACs (AP +1.6 mm, ML +0.6 mm, DV −6.6 mm) and received a fixed +0.8 V current via the auxiliary electrode. After insertion of the recording electrode, a series of 0.5 ms duration cathodal monophasic pulses (20 pulses at 50 Hz applied every 30 sec at 800 μA) were delivered to the stimulating electrode via an optical isolator and programmable pulse generator (Iso-Flex/Master-8; AMPI, Jerusalem, Israel) (Lester et al., 2010). Amperometry sessions began with a measure of phasic dopamine release evoked via electrical stimulation. Additionally, the synaptic half-life of dopamine following evoked release was quantified. The synaptic half-life of dopamine is defined as latency between peak dopamine release and restoration to 50% of baseline (Mittleman et al., 2011; Fielding et al., 2013). Next, subjects (*n* = 11) were injected with the dopamine transporter (DAT) inhibitor nomifensine (10 mg/kg, i.p.), while MFB stimulations continued every 30 seconds. We then measured dopamine half-life at 20, 40, and 60 minutes following nomifensine administration. The supply of available dopamine was measured by calculating total dopamine efflux during 3 minutes of continuous electrical stimulation with 9000 pulses at 50 Hz (Fielding et al., 2013). Following each amperometry experiment, an iron deposit was created to mark the stimulating electrode site by sending direct anodic current (100 μA for 10 s) through the electrode. Rats were then euthanized using a lethal intracardial injection of urethane (0.345 g/mL). Brains were removed and stored in a 30% sucrose / 10% formalin solution with 0.1% potassium ferricyanide for sectioning. Coronal sections of brains were sliced at −20°C using a cryostat, and electrode placements were identified using a light microscope and marked on coronal diagrams (Paxinos and Watson, 1997). *In vitro* electrode calibration was accomplished by exposing each carbon-fiber recording microelectrode to known solutions of dopamine (0.2 μM – 1.2 μM) via a flow injections system (Michael and Wightman, 1999; Prater et al., 2018).

### Statistical Analysis

Rats were separated into “Risk-Taking” and “Risk-Averse” groups as determined by a k-means cluster analysis on each subject’s percent choice of large risky reward in each block averaged across the final five days of training. This algorithm uses an iterative distance-from-center minimization technique to identify a user-specified number of clusters (MacQueen, 1967). High and low impulsivity groupings were divided “High-Impulsivity” and “Low-Impulsivity” via a similar criterion. Performance in the RDT and in DD was investigated using 2×5 repeated measures ANOVAs with risk or impulsivity group as a between subjects factor and % chance of shock or delay duration as a within subjects factor. Pearson correlations were used to test for linear relationships between risk preference and delay preference.

For amperometric recordings, dopamine efflux from .25s before to 10s after electrical stimulation was used to quantify baseline dopamine release, dopamine half-life, and sensitivity to DAT inhibiting drugs. Baseline (pre-drug) dopamine release was quantified in terms of peak extracellular dopamine concentrations following stimulation. Baseline dopamine half-life (calculated as time of return to baseline dopamine levels – time of peak dopamine levels / 2) served as a measure of DAT efficiency. The % change in dopamine half-life from pre-to post-drug provided a measure of susceptibility to the presence of a dopamine reuptake inhibitor (Mittleman et al., 2011; Fielding et al., 2013). Regional dopamine supply was quantified as the total concentration of dopamine release during a 3 minute continuous stimulation. Differences in baseline release, baseline half-life, and dopamine supply among risk and impulsivity groups were assessed using independent samples *t*-tests. Differences in post-drug dopamine half-life were examined using 2×3 repeated measures ANOVAs with risk or impulsivity group as a between subjects factor and time (20, 40, 60 min) as a within subjects factor. Pearson correlations were utilized to probe for linear relationships between risk-preference and measures of dopamine transmission, and were used to test for linear relationships between impulsive choice and dopamine release characteristics.

## Results

### Characterization of Risk Preference in the Risky Decision-Making Task

Analyses indicated a significant main effect of test block on the percentage of risky choices (*F*_(4, 72)_ = 19.14, *p* < 0.001, Figure 2 A). Thus, subjects showed a reduced preference for the risky reward as risk level increased. Risk-taking phenotypes were determined for each rat based on the average percentage of risky choices across sessions over the last 5 test days. A k-means clustering analysis was utilized to separate rats into groups based on risky decision-making variability: “risk-taking” rats, which demonstrated preference for the risky choice (*n* = 8), and “risk-averse” rats, which preferred the safe option (*n* = 11, Figure 2 B). Percentage of risky choices during test blocks 2 – 5 (when there was a probability of foot shock following risky choices) was statistically greater in risk-taking vs risk-averse groups as indicated by a main effect of risk group (*F*_(1, 17)_ = 136.07, *p* < 0.001); Figure 2 C – D). Although, no risk group x test block interaction was shown (*F*_(3, 51)_ = 0.27, *p* = 0.841).

**Figure 2.**
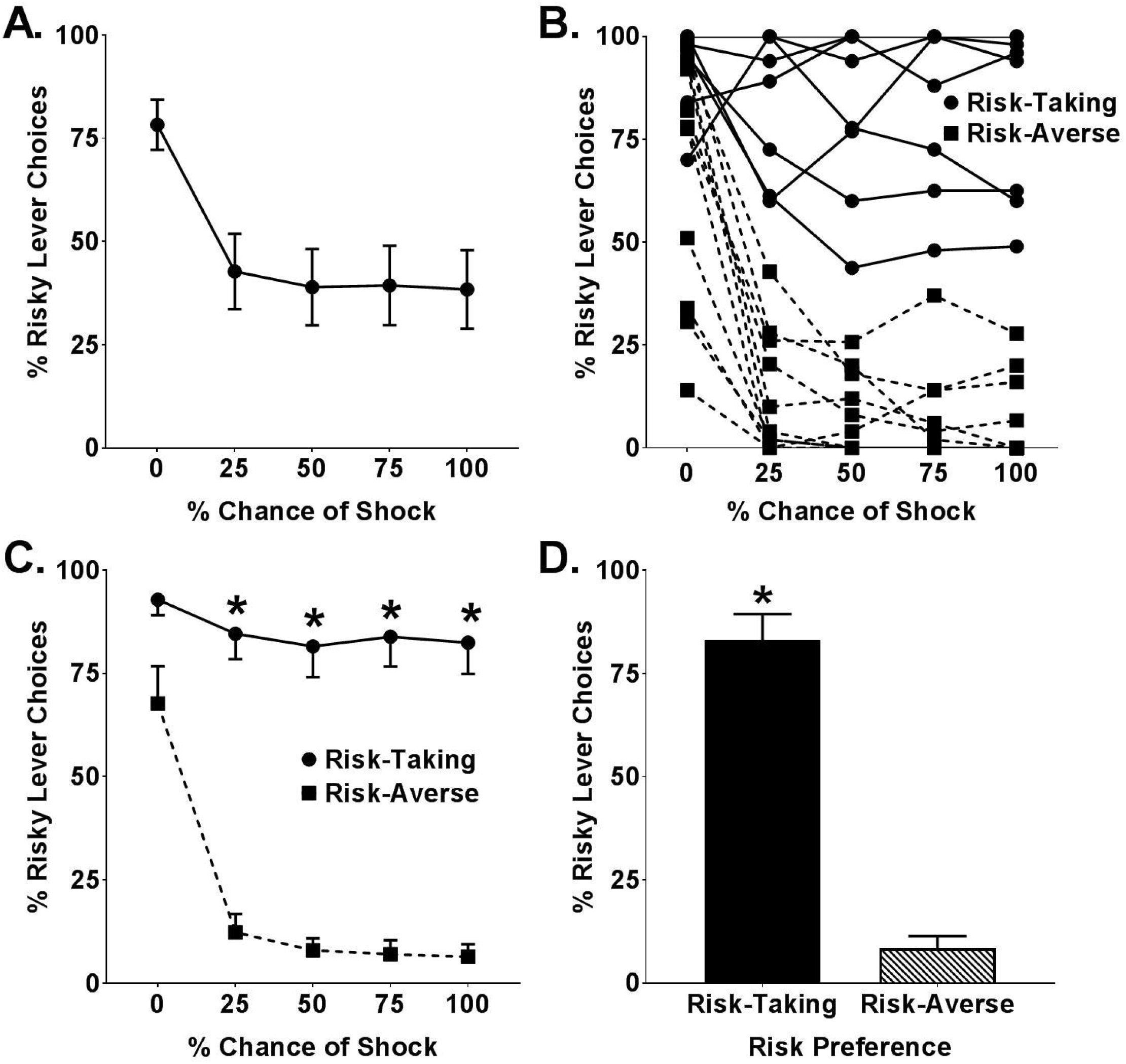
Performance on the risky decision-making task. *A*. Individual risk scores for each test block. *B*. Mean risk preference scores for each test block. *C*. Risk preference scores for entire test. Error bars represent ± *SEM*. \**p* < 0.05

### Assessment of Impulsive Choice Phenotypes in the Delay Discounting Task

Overall, a main effect of test block on percent delay choices indicated that rats shifted choice preference from large to small rewards as delay increased (*F*_(4, 72)_ = 48.44, *p* < 0.001, Figure 3 A). A k-means cluster analysis was again used to assign rats into low impulsive choice (*n* = 9) or high impulsive choice (*n* = 10, Figure 3 B) phenotypes. To validate impulsivity phenotype classifications, percent choice of delayed reward during blocks 2 – 5 were compared between the high and low impulsivity groups. Repeated measures analyses indicated a main effect of impulsivity group (*F*_(1, 17)_ = 33.91, *p* < 0.001) and a significant group by block interaction (*F*_(3, 51)_ = 3.94, *p* < 0.001; Figure 3 C – D).

**Figure 3.**
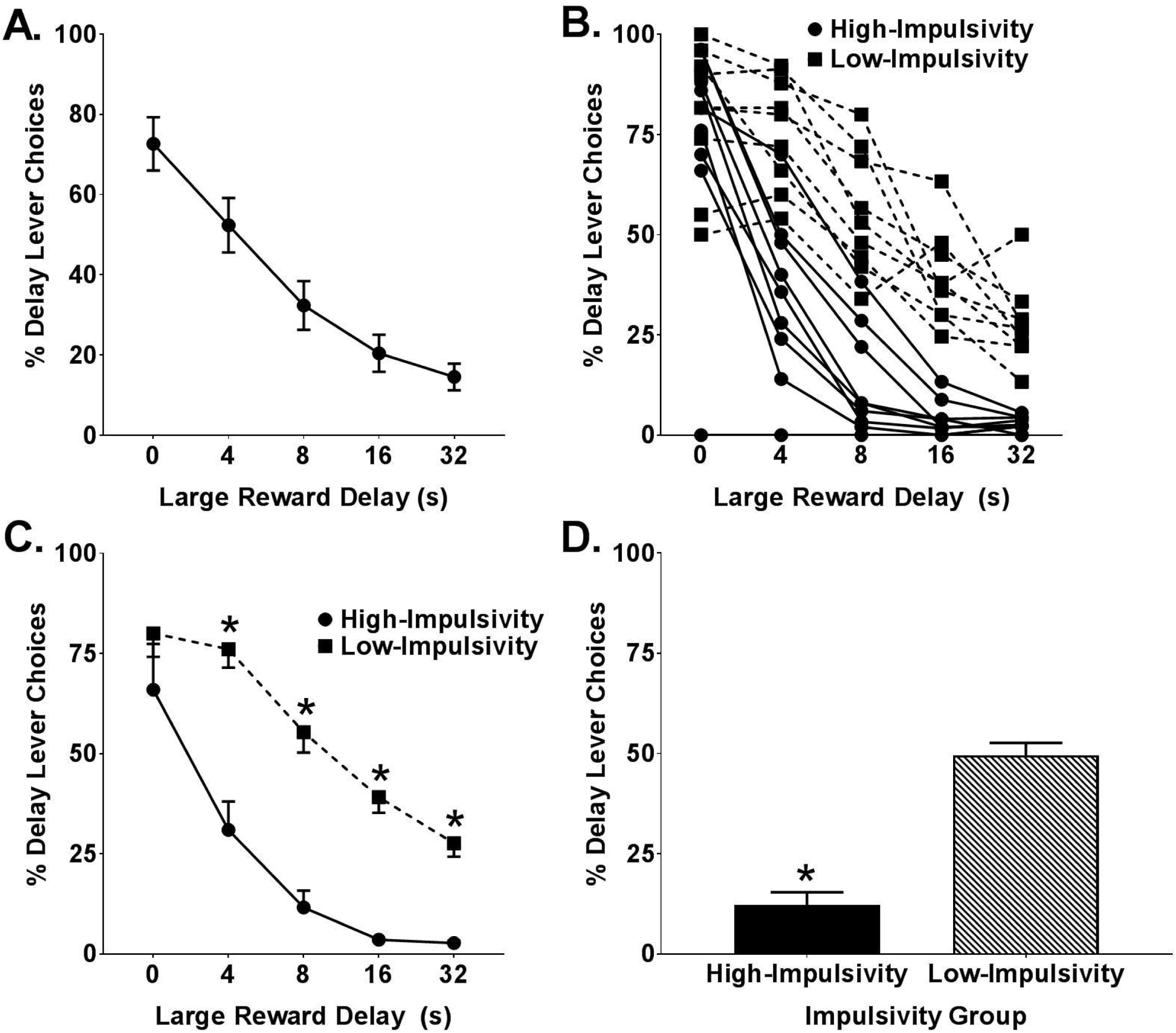
Performance on the delay discounting task. *A.* Individual impulsivity scores for each test block. *B.* Mean impulsivity scores for test blocks. *C.* Impulsivity scores across entire task. *D.* Scatter plot of total % delay choices by total % risky lever choices. Error bars represent ± *SEM*. *p < 0.05.

Next, we compared risky decision-making and delay discounting performance. There was no correlation between impulsive choice and risk-preference (*r* = 0.13, *p* = 0.578; Figure 4 A), suggesting that risk of punishment and delay produce different patterns of reward discounting. Moreover, we observed no difference between risk-takers and risk-averse in reward-preference during delay discounting (*F*_(1, 18)_ = 0.08, *p* = 0.775, Figure 4 B) and no difference between high-and low-impulsive rats on risk-taking on RDT (*F*_(1, 18)_ = 0.02, *p* = 0.888, Figure 4 C).

**Figure 4.**
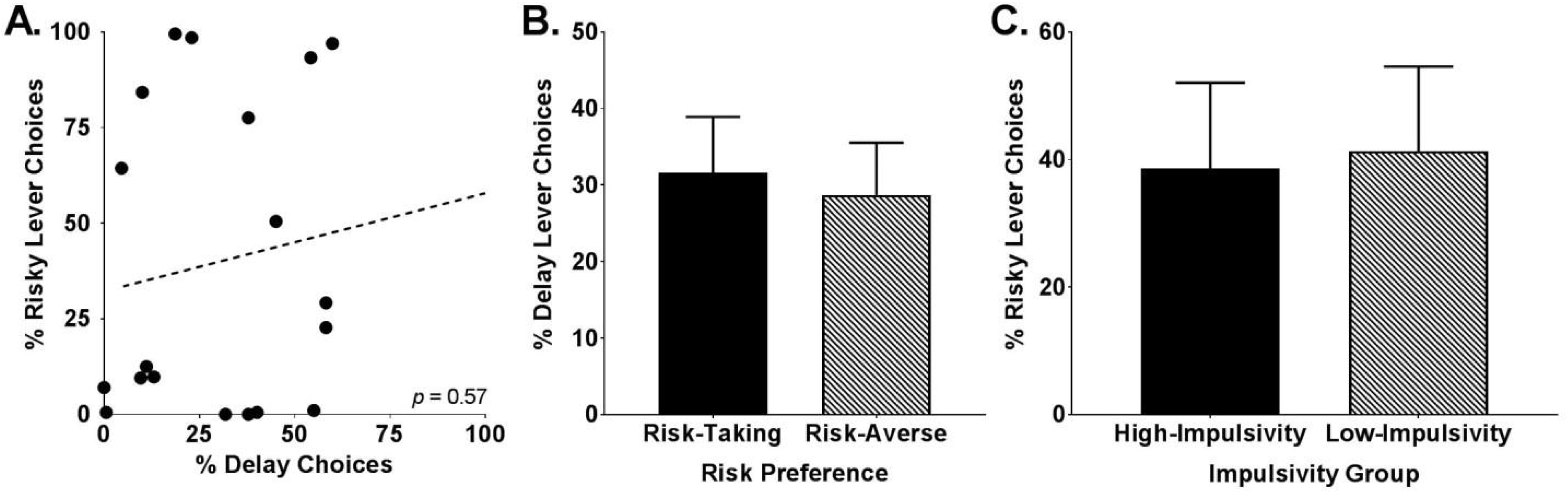
Comparison of risky decision-making and delay discounting performance. *A.* Correlation of scores from the risky decision-making and delay discounting tasks. *B.* Total percent delay lever choices for risk subpopulations. *C.* Total percent risky lever choices for impulsivity groups. Error bars represent ± *SEM.*

### Risk-Taking and Phasic Dopamine Dynamics in the Nucleus Accumbens

Fixed potential amperometry was used to characterize differences in dopamine release dynamics in the NACs of risk-taking and risk-averse rats. First, we measured MFB stimulation-evoked dopamine efflux in the NACs, which required accurate placement of electrodes (Figure 5). MFB stimulation successfully induced dopamine efflux in all subjects. Additionally, the range of baseline dopamine release in the NACs (0.62 ± 0.08 µM) and the percent increase in intra-NAC dopamine release following nomifensine administration (322.53 ± 60.48 %) were similar to previous studies (Freels et al., 2019; Holloway et al., 2019).

**Figure 5.**
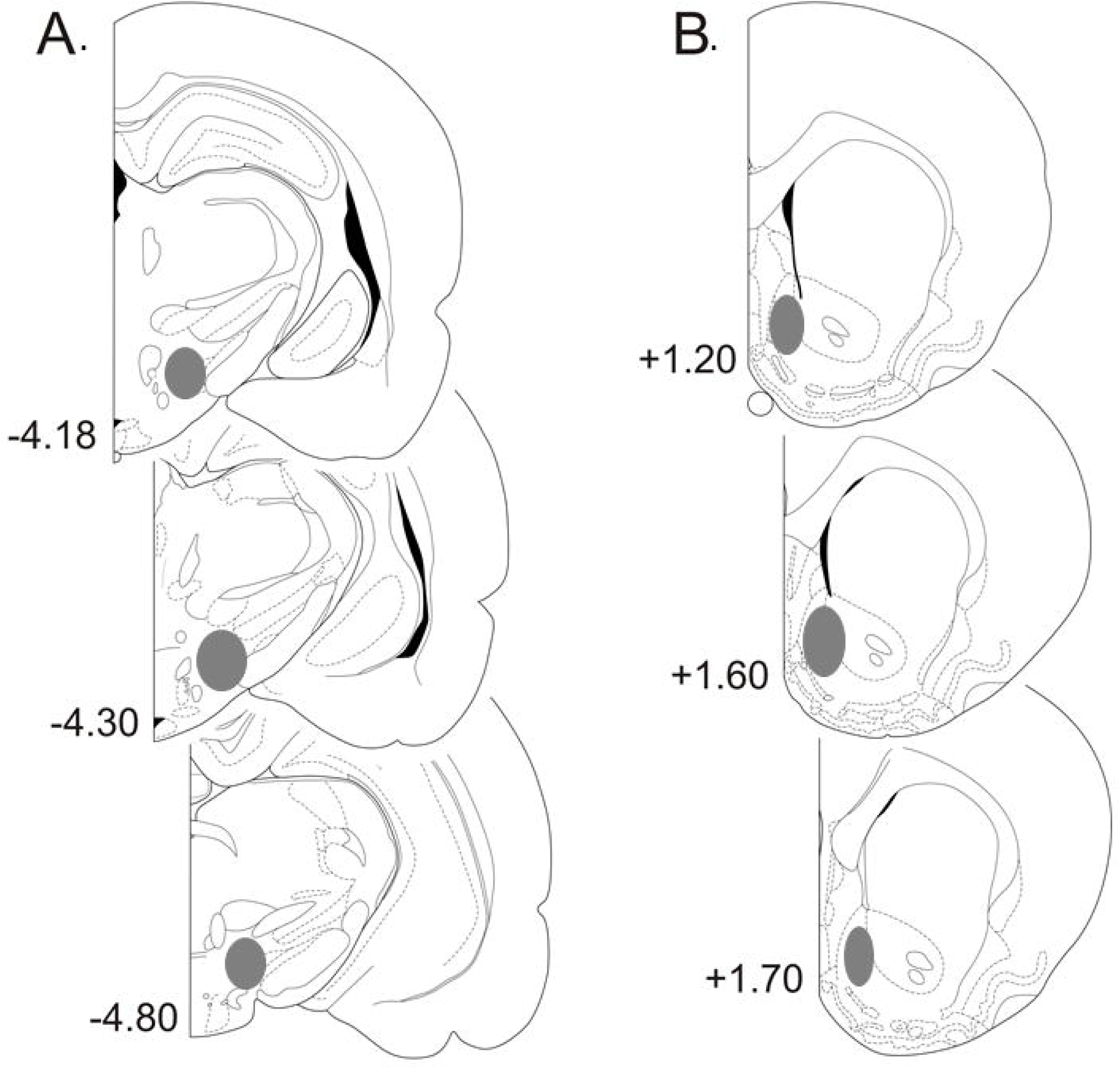
Representative coronal sections of the rat brain (adapted from the atlas of Paxinos and Watson, 2001), with grey shaded areas indicating the placements of (A) stimulating electrodes in the medial forebrain bundle and (B) amperometric recording electrodes in the nucleus accumbens. Numbers correspond to mm from bregma.

Analyses revealed significantly elevated dopamine release in risk-taking rats relative to risk averse rats (*t*_(17)_ = 3.85, *p* < .001; Figure 6 A – C). Pearson correlation analysis also revealed that risk preference was positively associated with evoked dopamine release (*r* = 0.54, *p* = .01; Figure 4 D). Dopamine supply was determined by continuously stimulating at 50 Hz for 3 min to evoke the release of all available dopamine in the NAC. Analyses indicated a trend toward increased dopamine supply in risk-taking relative to risk-averse rats (*t*_(9)_ = 1.99, *p* = .078; Figure 6 E). Additionally, there was a near significant positive correlation between risk-taking and dopamine supply (*r* = 0.56, *p* = .072, Figure 6 F).

**Figure 6.**
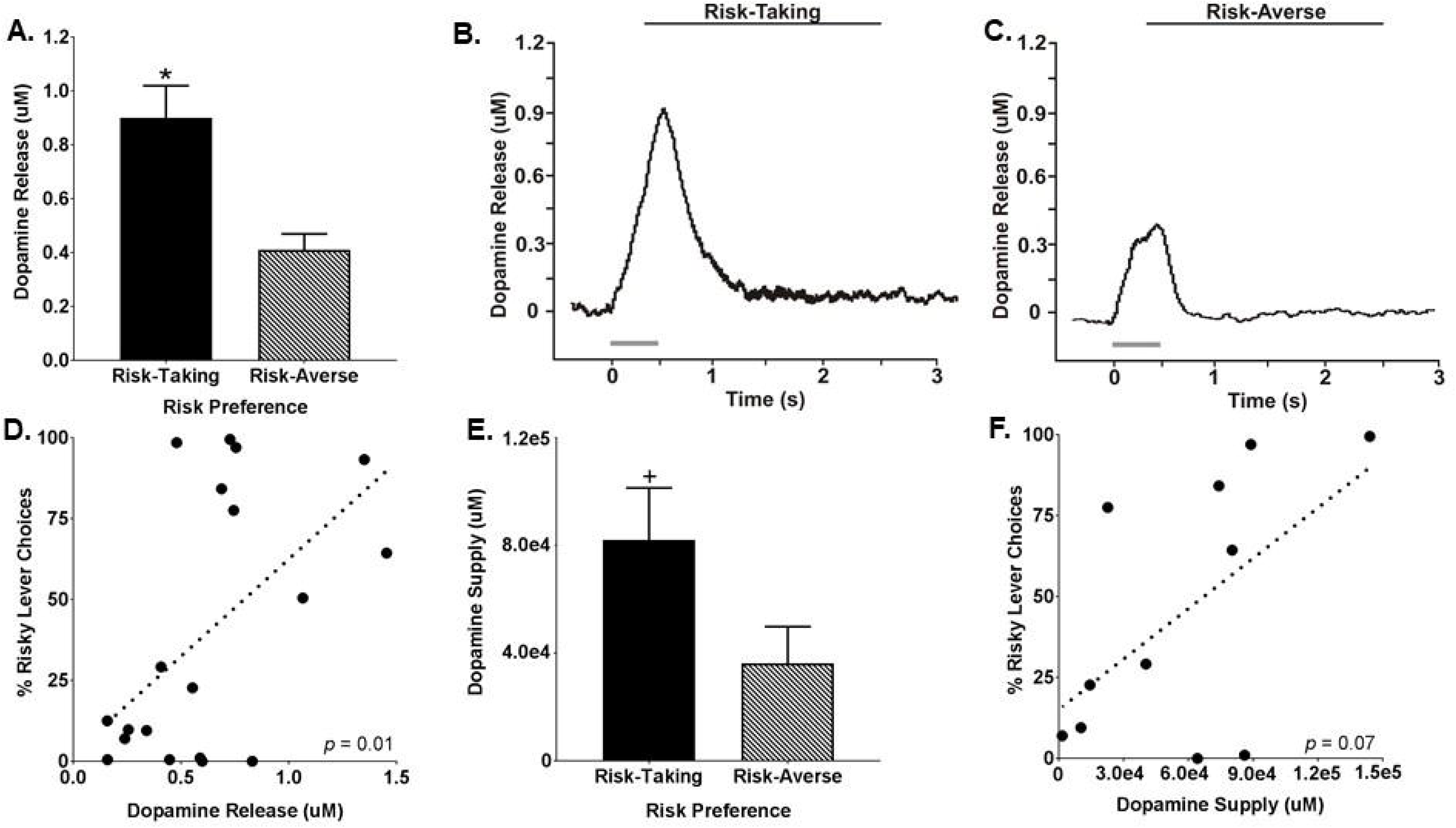
Baseline NACs dopamine release and dopamine supply in risk-taking and risk-averse subpopulations. *A.* Mean evoked dopamine release at baseline. *B.* Sample baseline response in risk-taking rats. *C.* Example baseline response in risk-averse rats. *D*. Mean dopamine supply in risk subpopulations. *E.* Correlation between total risk preference and dopamine release. *F*. Correlation between risk preference and dopamine supply. Error bars represent ± *SEM*. \**p* < 0.05. ^+^*p* = 0.07.

Next, we measured baseline dopamine half-life as a function of risk preference. On average, risk-taking and risk-averse rats demonstrate similar baseline durations of dopamine half-life (*t*_(17)_ = −0.89, *p* = .387; Figure 7A). We then assessed sensitivity to the DAT inhibitor nomifensine in risk-taking and risk-averse phenotypes. This was determined by measuring the % change in dopamine half-life at 20 minute intervals post systemic nomifensine administration for 1 hour. Repeated measures analyses revealed a significant risk group x time point interaction on % change in dopamine half-life following nomifensine administration (*F*_(3, 27)_ = 2.99, *p* = .048; Figure 5B). Subsequent analyses revealed that dopamine half-life was increased in risk-taking rats at the 40 (*t*_(9)_ = 2.60, *p* = .029) and 60 minute time points (*t*_(9)_ = 2.65, *p* = .027) relative to risk averse rats (Figure 7B). Additionally, correlation analyses showed significant positive relationships between risk preference and the average percent change in dopamine half-life across the 20, 40, and 60 minute time points (*r* = 0.63, *p* = .035; Figure 7C). In summary, dopamine half-life is comparable between risk-taking and risk-averse rats at baseline, but is significantly elevated in risk-taking rats following exposure to a DAT inhibitor.

**Figure 7.**
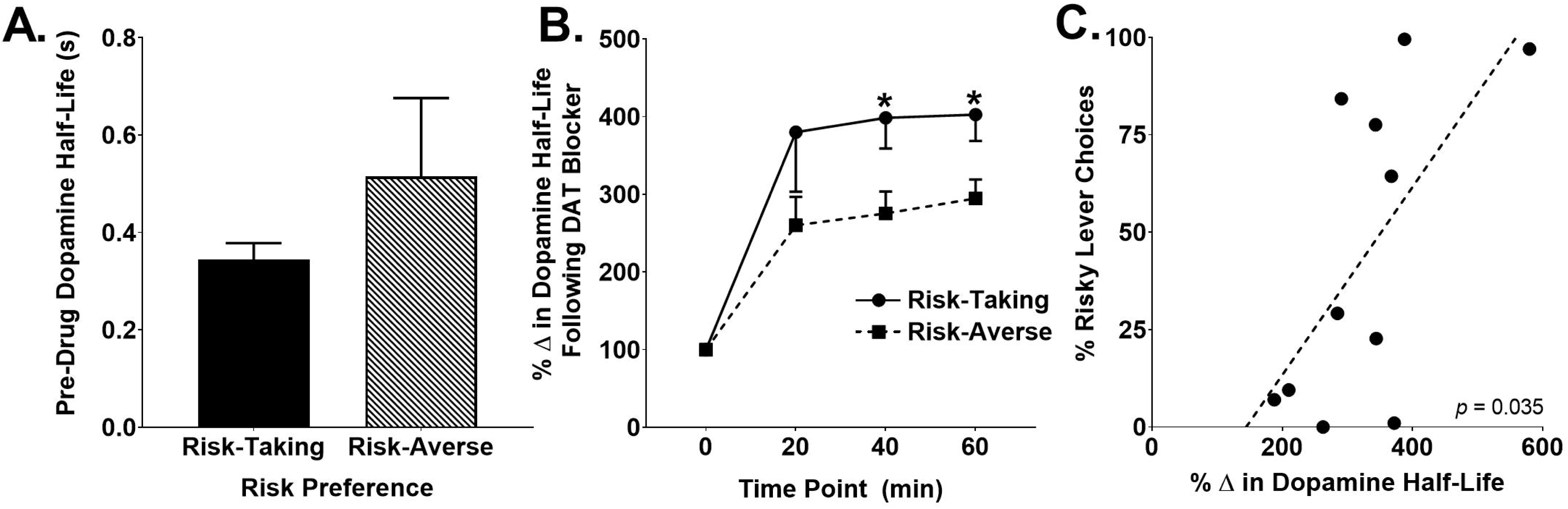
Dopamine half-life measures for risk-taking and risk-averse phenotypes. *A*. Baseline dopamine half-life for risk subpopulations. *B*. Percent change in dopamine half-life following nomifensine treatment during 60 minute amperometric recordings for risk-taking and -averse rats. *C*. Correlation between risk preference and average percent change in dopamine half-life across the 20, 40, and 60 minute time points. Error bars represent ± *SEM*. \**p* < 0.05.

### Delay Discounting and Phasic Dopamine Dynamics in the Nucleus Accumbens

Phasic dopamine release dynamics in the NACs was also compared to impulsive choice, with subjects divided into high and low impulsive responders. Analyses indicated no significant main effect of impulsivity group on evoked dopamine release (*t*_(17)_ = −1.03, *p* = .317; Figure 8A-C), and no correlation between dopamine release in the NACs and percent delay lever choices (*r* = 0.23, *p* = .349, Figure 8D). There was also no main effect of impulsivity group on overall dopamine supply in NACs (*t*_(9)_ = 0.43, *p* = 0.670; Figure 8E). Moreover, correlation analyses revealed that dopamine supply and percent delay lever choices are not related (*r* = 0.04, *p* = .899, Figure 8F).

**Figure 8.**
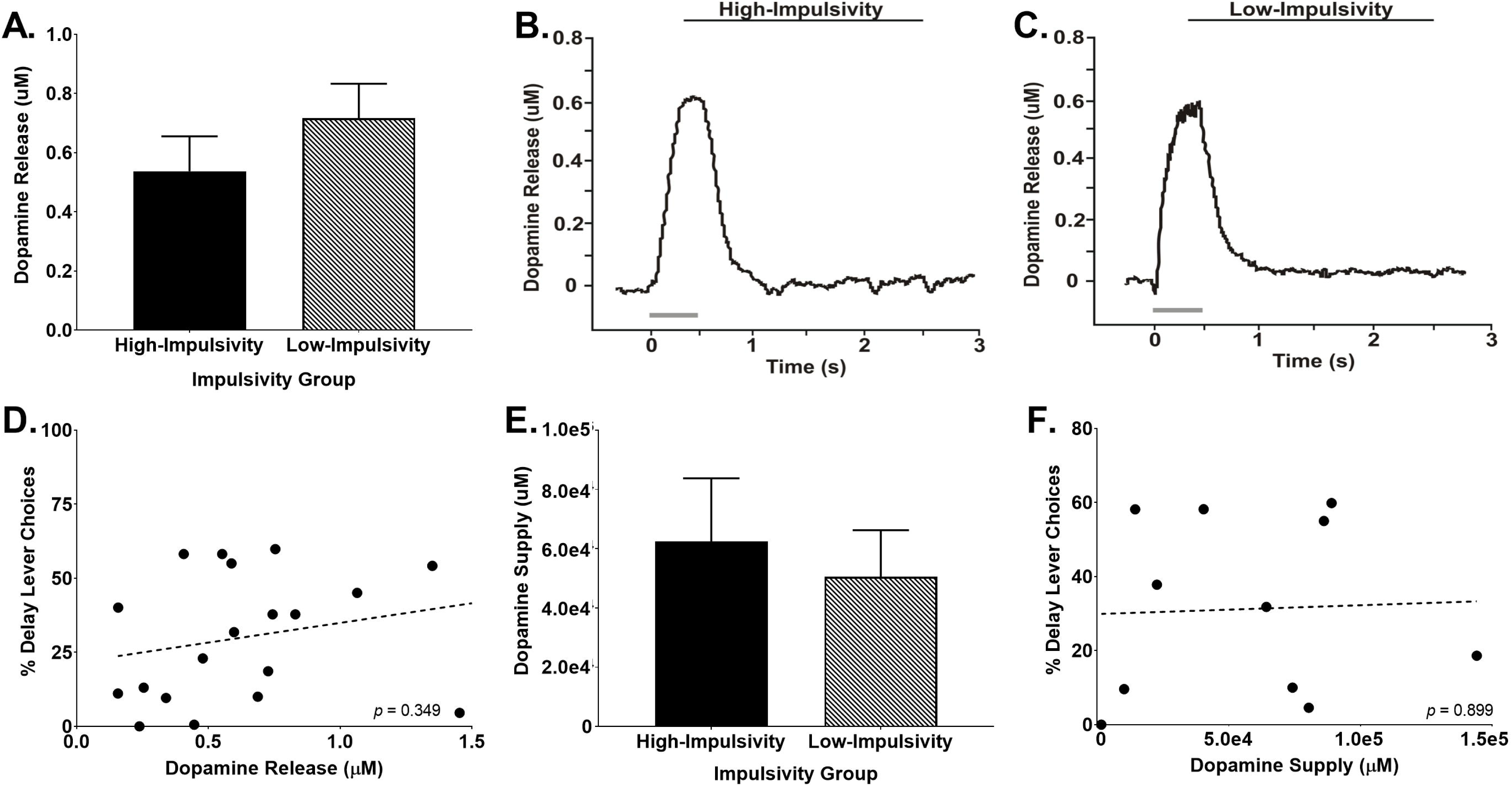
Baseline evoked dopamine release and dopamine supply for impulsivity subpopulations. *A.* Mean evoked dopamine release at baseline. *B.* Sample baseline response in high impulsive choice rats. *C.* Example baseline response in low impulsive choice rats. *D*. Mean dopamine supply in impulsivity phenotypes. *E.* Correlation between baseline dopamine release and total delay preference. *F*. Correlation between dopamine supply and impulsive choice. Error bars represent ± *SEM*.

## Discussion

Here, we utilized fixed potential amperometry to examine evoked dopamine release in the NACs of anesthetized rats previously characterized in both the RDT and delay discounting. Consistent with previous studies, we observed no relationship between punishment-based risky decision-making and impulsive choice (Simon et al., 2009; Gabriel et al., 2018). MFB stimulation-evoked dopamine release in NACs was both positively correlated with risk preference and enhanced in risk-taking compared to risk-averse rats. There was also a trend toward enhanced dopamine supply in risk-takers, and risk preference predicted sensitivity to the DAT blocker nomifensine, quantified as increased dopamine half-life in the synapse after MFB stimulation. Conversely, evoked dopamine release and dopamine supply were not predicted by individual differences in impulsive choice.

### Risky Decision-making predicts phasic dopamine release in NAC

Risk-taking in the RDT has been associated with dopamine receptor expression in NACs ex vivo (Simon et al., 2011; Mitchell et al., 2014), yet little is known about the relationship between risk-taking and biologically relevant dopamine signaling. The current observation of elevated NACs phasic dopamine release suggests that risk-taking rats may be more susceptible to the motivational effects of environmental cues, as phasic dopamine release mediates the attribution of motivational salience to reward-predictive stimuli (Flagel et al., 2011). This is consistent with risk-taking predicting enhanced salience attributed to reward-predictive cues, manifested as elevated sign-tracking (Olshavsky et al., 2014). These data also suggest that the previously observed abundance of D1 receptors in the NACs of risk-taking rats may be a response to enhanced dopamine activity in the region (Simon et al., 2011). This exaggerated dopamine response may be the impetus for persistent seeking of rewards in the face of punishment.

The magnitude of a stimulation-evoked dopamine response is in part determined by the rate of dopamine reuptake. However, given that there were no differences between risk groups in the synaptic half-life of dopamine, the observed differences in dopamine release is likely not related to DAT functioning. Dopamine half-life is related to DAT efficiency, with a shorter half-life being indicative of greater efficiency (Benoit-Marand et al., 2000; Mittleman et al., 2011), suggesting that reuptake following dopamine release does not vary as a function of risk-taking phenotype. In addition, we observed a near-significant trend toward enhanced dopamine supply in risk-taking rats. This may be a compensatory mechanism resulting from elevated phasic dopamine release. Conversely, elevated dopamine supply may contribute to enhanced phasic dopamine release in risk takers; further experimentation is necessary to confirm the causal direction of this relationship.

Risky decision-making is predictive of impulsive action (Gabriel et al., 2018), a facet of impulsivity distinct from impulsive choice characterized by ongoing reward-seeking despite unfavorable outcomes (Everitt & Robbins, 2016; Dalley & Robbins, 2017). Like risk-taking, impulsive action is mediated through striatal dopamine activity (Robbins, 2002; Pattij et al., 2007; Besson et al., 2010; Mitchell et al., 2014; Zalocusky et al., 2016). In addition, both impulsive action and risk-taking predict cocaine self-administration (Dalley et al., 2007; Belin et al., 2008; Mitchell et al., 2014). Thus, elevated phasic dopamine release in NACs may produce the co-expression of both impulsive and risk-taking phenotypes, in addition to increased vulnerability to psychostimulant drugs of abuse. This cluster of addiction-relevant behavioral traits in conjunction with elevated dopamine sensitivity suggests that the RDT may have utility for detecting both cognitive and neuropharmacological vulnerability to drugs of abuse.

Previous studies have examined the relationship between dopamine in NAC and probabilistic discounting, which quantifies decision-making governed by the risk of reward omission rather than punishment (Orsini et al., 2015a). Tonic dopamine efflux in NAC has been shown via microdialysis to vary at the same rate as preference for increasingly probabilistic rewards (St. Onge et al., 2012). In addition, dopamine neuron activity scales with probability of reward delivery (Fiorillo et al., 2003). While this provides evidence for dopamine’s role in behavioral flexibility in the face of uncertain rewards, it does not take into account the relationship between individual differences in risky decision-making and dopamine release dynamics. Sugam et al (2012) observed that NAC dopamine release increases in response to cues predicting rewards of subjectively greater value during probabilistic discounting, suggesting that phasic dopamine release during risk-taking is associated with baseline risk-preference (Sugam et al., 2012). The current data expand upon this by demonstrating that differences in dopamine sensitivity are not limited to risk-associated events, but can be observed after general axonal stimulation. Thus, elevated phasic dopamine release in risk-takers likely manifests in response to a range of situations that elicit dopamine neuron firing, potentially causing vulnerability to pathologies of motivation.

Another critical distinction between this experiment and other studies is operationalization of risky decision-making. Probability discounting, utilized by (St. Onge and Floresco, 2010; Sugam et al., 2012; Zalocusky et al., 2016), defines risk as probability of reward omission, whereas RDT utilizes probability of punishment and is designed to model situations in which rewards are consistently delivered, but there is a risk of a negative outcome that differs in modality from the reward (Orsini et al., 2015a). This punishment-driven risk taking can be more readily extrapolated to the risk-taking performed during SUDs, in which the reward (drug reinforcement) often differs from the consequences (risk of arrest or overdose). The distinction between these forms of risk-taking is further supported by divergent effects of pharmacological and neurobiological manipulations (St. Onge & Floresco, 2009; Simon et al., 2011; Stopper et al., 2014; Orsini et al., 2015a, 2015b). Therefore, these data expand upon previous studies by being the first to characterize the relationship between punishment-based risk-taking and NAC dopamine release dynamics, in addition to the first to selectively examine dopamine release in the shell region.

### Risk-taking predicts sensitivity to dopamine transporter blockade

Risk-taking rats demonstrated elevated sensitivity to the DAT inhibitor nomifensine, reflected as increased latency for dopamine to restore to baseline levels after phasic activation. This may be another indicator of vulnerability to drugs of abuse in the risk-taking subpopulation. Elevated sensitivity to nomifensine translates to increased sensitivity to dopaminergic drugs of abuse that affect DAT, which include cocaine and amphetamine (Carboni et al., 1989; Kuczenski and Segal, 1989; Mcelvain & Schenk, 1992). Consistent with this finding, risk-taking rats have previously been shown to self-administer more cocaine than risk-averse rats (Mitchell et al., 2014). A possible explanation for increased sensitivity to nomifensine is diminished DAT capacity (Mittleman et al., 2011). However, this is unlikely due to the similarities in dopamine half-life prior to nomifensine. A more plausible explanation is that elevated post-nomifensine half-life is a function of elevated dopamine supply causing a greater concentration of dopamine to flood the synapse after stimulation, requiring an extended time for clearance.

### Delay discounting is not associated with phasic dopamine release

We found no relationship between phasic dopamine release and delay discounting, a measure of preference for immediate vs delayed rewards. This was somewhat surprising, as past research has found dopamine to be involved with modulation of impulsive choice, with dopamine receptor activation decreasing impulsivity (Wade et al., 2000; van Gaalen et al., 2006). However, NAC dopamine depletion has no effect on delay discounting (Winstanley et al., 2005), which is consistent with our observation that dopamine supply does not predict impulsive choice. Furthermore, a recent report showed associations between delay discounting and dopamine receptors were restricted to clinical populations, with no relationship with healthy individuals (Castrellon et al., 2019). Therefore, while dopamine may play a modulatory role in delay discounting, it is not a reliable correlate of individual differences in impulsive choice in non-pathological populations. This demonstrates that elevated dopamine release is specific to risky decision-making and does not generalize to all forms of economic decision-making.

## Conclusions

Risky decision-making is commonly observed in SUD (Bechara et al., 2001; Brand et al., 2008; Brevers et al., 2014; Verdejo-Garcia et al., 2018). Therefore, understanding the neurobiology underlying specific biases in risk-taking may have utility for early detection and treatment of vulnerable individuals. These data reveal multiple measures of a hyper-sensitive mesolimbic dopamine system in rats predisposed to risk-taking, while demonstrating a clear distinction in the dopaminergic correlates of these two forms of decision-making (risk-taking vs delay discounting). This suggests NACs phasic dopamine dynamics as a potential therapeutic target for pathological risk-taking, but not impulsive choice. This preclinical identification of enhanced dopamine sensitivity in a readily identifiable subpopulation of rats may provide further utility in additional studies investigating the neuronal and genetic correlates of vulnerability to SUD, as well as in the identification of therapeutic targets for prevention and treatment.

## Acknowledgements

We would like to thank Amber Woods, Haleigh Joyner, Samantha Morrison, Andrew Starnes, and Alan Rasheed for technical assistance. This research was supported by a Young Investigator Grant from the Brain and Behavior Research Foundation and a Faculty Research Grant from the University of Memphis (NWS).

## References

Andrade LF, Petry NM (2012) Delay and probability discounting in pathological gamblers with and without a history of substance use problems. Psychopharmacology (Berl) 219:491–499.

Barlow RL, Gorges M, Wearn A, Niessen HG, Kassubek J, Dalley JW, Pekcec A (2018) Ventral Striatal D2/3 Receptor Availability Is Associated with Impulsive Choice Behavior As Well As Limbic Corticostriatal Connectivity. Int J Neuropsychopharmacol 21:705–715 Available at: https://academic.oup.com/ijnp/article/21/7/705/4938500.

Bechara A, Dolan S, Denburg N, Hindes A, Anderson SW, Nathan PE (2001) Decision-making deficits, linked to a dysfunctional ventromedial prefrontal cortex, revealed in alcohol and stimulant abusers. Neuropsychologia 39:376–389.

Belin D, Mar AC, Dalley JW, Robbins TW, Everitt BJ (2008) High Impulsivity Predicts the Switch to Compulsive Cocaine-Taking. Science (80-) 320:1352–1356.

Benoit-Marand M, Jaber M, Gonon F (2000) Release and elimination of dopamine in vivo in mice lacking the dopamine transporter: Functional consequences. Eur J Neurosci 12:2985– 2992.

Benoit-Marand M, Suaud-Chagny M-F, Gonon F (2007) Presynaptic Regulation of Extracellular Dopamine as Studied by Continuous Amperometry in Anesthetized Animals. In: Electrochemical Methods for Neuroscience (Michael AC, Borland LM, eds). Frontiers in Neuroengineering.

Besson M, Belin D, McNamara R, Theobald DEH, Castel A, Beckett VL, Crittenden BM, Newman AH, Everitt BJ, Robbins TW, Dalley JW (2010) Dissociable control of impulsivity in rats by dopamine D2/3 receptors in the core and shell subregions of the nucleus accumbens. Neuropsychopharmacology 35:560–569 Available at: http://dx.doi.org/10.1038/npp.2009.162.

Bickel WK, Johnson MW, Koffarnus MN, MacKillop J, Murphy JG (2014) The Behavioral Economics of Substance Use Disorders: Reinforcement Pathologies and Their Repair. Annu Rev Clin Psychol 10:641–677 Available at: http://www.annualreviews.org/doi/10.1146/annurev-clinpsy-032813-153724.

Brand M, Roth-Bauer M, Driessen M, Markowitsch HJ (2008) Executive functions and risky decision-making in patients with opiate dependence. Drug Alcohol Depend 97:64–72.

Brevers D, Bechara A, Cleeremans A, Kornreich C, Verbanck P, Noël X (2014) Impaired decision-making under risk in individuals with alcohol dependence. Alcohol Clin Exp Res 38:1924–1931.

Caitlin A. Orsini, Markie L. Willis, Ryan J. Gilbert, Jennifer L. Bizon BS, Orsini CA, Willis ML, Gilbert RJ, Bizon JL, Setlow B (2016) Sex Differences in a Rat Model of Risky Decision Making. Behav Neurosci 130:1922–2013.

Carboni E, Imperato A, Perezzani L, Di Chiara G (1989) Amphetamine, cocaine, phencyclidine and nomifensine increase extracellular dopamine concentrations preferentially in the nucleus accumbens of freely moving rats. Neuroscience 28:653–661.

Castrellon JJ, Seaman KL, Crawford JL, Young JS, Smith CT, Dang LC, Hsu M, Cowan RL, Zald DH, Samanez-Larkin GR (2019) Individual differences in dopamine are associated with reward discounting in clinical groups but not in healthy adults. J Neurosci 39:321–332.

Coffey SF, Gudleski GD, Saladin ME, Brady KT (2003) Impulsivity and rapid discounting of delayed hypothetical rewards in cocaine-dependent individuals. Exp Clin Psychopharmacol.

Dalley JW, Fryer TD, Brichard L, Robinson ESJ, Theobald DEH, Laane K, Pena Y, Murphy ER, Shah Y, Probst K, Abakumova I, Aigbirhio FI, Richards HK, Hong Y, Baron J-C, Everitt BJ, Robbins TW (2007) Nucleus Accumbens D2/3 Receptors Predict Trait Impulsivity and Cocaine Reinforcement. Science (80-) 315:1267–1270 Available at: http://www.sciencemag.org/cgi/doi/10.1126/science.1137073.

Dalley JW, Robbins TW (2017) Fractionating impulsivity: neuropsychiatric implications. Nat Rev Neurosci 18:158–171 Available at: http://www.nature.com/doifinder/10.1038/nrn.2017.8.

Dugast C, Suaud-Chagny MF, Gonon F (1994) Continuous in vivo monitoring of evoked dopamine release in the rat nucleus accumbens by amperometry. Neuroscience.

Everitt BJ, Robbins TW (2016) Drug Addiction: Updating Actions to Habits to Compulsions Ten Years On. Annu Rev Psychol 67:23–50 Available at: http://www.annualreviews.org/doi/10.1146/annurev-psych-122414-033457.

Fielding JR, Rogers TD, Meyer AE, Miller MM, Nelms JL, Mittleman G, Blaha CD, Sable HJK (2013) Stimulation-evoked dopamine release in the nucleus accumbens following cocaine administration in rats perinatally exposed to polychlorinated biphenyls. Toxicol Sci 136:144–153.

Fiorillo CD, Tobler PN, Schultz W (2003) Discrete Coding of Reward Dopamine Neurons. Science (80-) 299:1898–1902 Available at: http://www.sciencemag.org/content/299/5614/1898.short.

Flagel SB, Clark JJ, Robinson TE, Mayo L, Czuj A, Willuhn I, Akers CA, Clinton SM, Phillips PEM, Akil H (2011) A selective role for dopamine in stimulus-reward learning. Nature 469:53–59 Available at: http://dx.doi.org/10.1038/nature09588.

Floresco SB, West AR, Ash B, Moorel H, Grace AA, Moore H, Grace AA, Moorel H, Grace AA (2003) Afferent modulation of dopamine neuron firing differentially regulates tonic and phasic dopamine transmission. Nat Neurosci 6:968–973 Available at: http://www.nature.com/doifinder/10.1038/nn1103.

Freels TG, Lester DB, Cook MN (2019) Arachidonoyl serotonin (AA-5-HT) modulates general fear-like behavior and inhibits mesolimbic dopamine release. Behav Brain Res 362:140– 151.

Gabriel DBK, Freels TG, Setlow B, Simon NW (2018) Risky decision-making is associated with impulsive action and sensitivity to first-time nicotine exposure. Behav Brain Res:0–1 Available at: https://www.sciencedirect.com/science/article/pii/S0166432818310118?via%3Dihub.

Gabriel DBK, Freels TG, Setlow B, Simon NW (2019) Risky decision-making is associated with impulsive action and sensitivity to first-time nicotine exposure. Behav Brain Res 359.

Garavan H, Hester R (2007) The role of cognitive control in cocaine dependence. Neuropsychol Rev 17:337–345.

Goldstein RZ, Volkow ND (2002) Drug addiction and its underlying neurobiological basis: Neuroimaging evidence for the involvement of the frontal cortex. Am J Psychiatry.

Heil SH, Johnson MW, Higgins ST, Bickel WK (2006) Delay discounting in currently using and currently abstinent cocaine-dependent outpatients and non-drug-using matched controls. Addict Behav.

Holloway ZR, Freels TG, Comstock JF, Nolen HG, Sable HJ, Lester DB (2019) Comparing phasic dopamine dynamics in the striatum, nucleus accumbens, amygdala, and medial prefrontal cortex. Synapse 73:e22074.

Ikemoto S, Panksepp J (1999) The role of nucleus accumbens dopamine in motivated behavior: A unifying interpretation with special reference to reward-seeking. Brain Res Rev 31:6–41.

Kuczenski R, Segal D (1989) Concomitant characterization of behavioral and striatal neurotransmitter response to amphetamine using in vivo microdialysis. J Neurosci 9:2051– 2065.

Lester DB, Miller AD, Blaha CD (2010) Muscarinic receptor blockade in the ventral tegmental area attenuates cocaine enhancement of laterodorsal tegmentum stimulation-evoked accumbens dopamine efflux in the mouse. Synapse.

MacQueen J (1967) Some Methods for classification and Analysis of Multivariate Observations. In: 5th Berkeley Symposium on Mathematical Statistics and Probability 1967.

Mcelvain JS, Schenk JO (1992) A multisubstrate mechanism of striatal dopamine uptake and its inhibition by cocaine. Biochem Pharmacol 43:2189–2199.

Michael DJ, Wightman RM (1999) Electrochemical monitoring of biogenic amine neurotransmission in real time. J Pharm Biomed Anal.

Mitchell MR, Vokes CM, Blankenship AL, Simon NW, Setlow B (2011) Effects of acute administration of nicotine, amphetamine, diazepam, morphine, and ethanol on risky decision-making in rats. Psychopharmacology (Berl) 218:703–712.

Mitchell MR, Weiss VG, Beas BS, Morgan D, Bizon JL, Setlow B (2014) Adolescent risk taking, cocaine self-administration, and striatal dopamine signaling. Neuropsychopharmacology 39:955–962 Available at: http://www.ncbi.nlm.nih.gov/pubmed/24145852.

Mittleman G, Call SB, Cockroft JL, Goldowitz D, Matthews DB, Blaha CD (2011) Dopamine dynamics associated with, and resulting from, schedule-induced alcohol self-administration: Analyses in dopamine transporter knockout mice. Alcohol 45:325–339.

Mogenson GJ, Jones DL, Yim Chiyiu (1980) FROM MOTIVATION TO ACTION?: FUNCTIONAL INTERFACE BETWEEN THE LIMBIC SYSTEM AND THE MOTOR SYSTEM inte. Animals 14:69–97.

Olshavsky ME, Shumake J, Rosenthal AA, Kaddour-Djebbar A, Gonzalez-Lima F, Setlow B, Lee HJ (2014) Impulsivity, risk-taking, and distractibility in rats exhibiting robust conditioned orienting behaviors. J Exp Anal Behav 102:162–178.

Orsini CA, Moorman DE, Young JW, Setlow B, Floresco SB (2015a) Neural mechanisms regulating different forms of risk-related decision-making: Insights from animal models. Neurosci Biobehav Rev 58:147–167 Available at: http://dx.doi.org/10.1016/j.neubiorev.2015.04.009.

Orsini CA, Trotta RT, Bizon JL, Setlow B (2015b) Dissociable Roles for the Basolateral Amygdala and Orbitofrontal Cortex in Decision-Making under Risk of Punishment. J Neurosci 35:1368–1379.

Pattij T, Janssen MCW, Vanderschuren LJMJ, Schoffelmeer ANM, van Gaalen MM (2007) Involvement of dopamine D1 and D2 receptors in the nucleus accumbens core and shell in inhibitory response control. Psychopharmacology (Berl) 191:587–598 Available at: http://link.springer.com/10.1007/s00213-006-0533-x.

Paxinos G, Watson C (1997) The Rat Brain in Stereotaxic Coordinates. Acad Press San Diego 3rd.

Perry JL, Carroll ME (2008) The role of impulsive behavior in drug abuse. Psychopharmacology (Berl) 200:1–26.

Prater WT, Swamy M, Beane MD, Lester DB (2018) Examining the Effects of Common Laboratory Methods on the Sensitivity of Carbon Fiber Electrodes in Amperometric Recordings of Dopamine. J Behav Brain Sci 8:117–125 Available at: http://www.scirp.org/journal/jbbs.

Robbins TW (2002) The 5-choice serial reaction time task: behavioural pharmacology and functional neurochemistry. Psychopharmacology (Berl) 163:362–380.

Robinson TE, Berridge KC (1993) The neural basis of drug craving: an incentive salience theory of addiction. Brain Res Rev 8:247–291 Available at: http://www.ncbi.nlm.nih.gov/pubmed/8401595%5Cnhttp://scholar.google.com/scholar?hl=en&btnG=Search&q=intitle:The+neural+basis+of+drug+craving+:+an+incentive+salience+theory#3.

Saddoris MP, Sugam JA, Stuber GD, Witten IB, Deisseroth K, Carelli RM (2015) Mesolimbic Dopamine Dynamically Tracks, and Is Causally Linked to, Discrete Aspects of Value-Based Decision Making. Biol Psychiatry 77:903–911 Available at: http://dx.doi.org/10.1016/j.biopsych.2014.10.024.

Setlow B, Mendez IA, Mitchell MR, Simon NW (2009) Effects of chronic administration of drugs of abuse on impulsive choice (delay discounting) in animal models. Behav Pharmacol.

Simon NW, Gilbert RJ, Mayse JD, Bizon JL, Setlow B (2009) Balancing Risk and Reward: A Rat Model of Risky Decision Making. Neuropsychopharmacology 34:2208–2217 Available at: http://www.nature.com/doifinder/10.1038/npp.2009.48.

Simon NW, Mendez IA, Setlow B (2007) Cocaine exposure causes long-term increases in impulsive choice. Behav Neurosci 121:543–549 Available at: http://doi.apa.org/getdoi.cfm?doi=10.1037/0735-7044.121.3.543.

Simon NW, Montgomery KS, Beas BS, Mitchell MR, LaSarge CL, Mendez IA, Banuelos C, Vokes CM, Taylor AB, Haberman RP, Bizon JL, Setlow B (2011) Dopaminergic Modulation of Risky Decision-Making. J Neurosci 31:17460–17470 Available at: http://www.jneurosci.org/cgi/doi/10.1523/JNEUROSCI.3772-11.2011.

Simon NW, Setlow B (2012) Modeling Risky Decision Making in Rodents. Methods Mol Biol 829:165–175 Available at: http://link.springer.com/10.1007/978-1-61779-458-2.

Sombers LA, Beyene M, Carelli RM, Mark Wightman R (2009) Synaptic Overflow of Dopamine in the Nucleus Accumbens Arises from Neuronal Activity in the Ventral Tegmental Area. J Neurosci.

St. Onge JR, Ahn S, Phillips AG, Floresco SB (2012) Dynamic Fluctuations in Dopamine Efflux in the Prefrontal Cortex and Nucleus Accumbens during Risk-Based Decision Making. J Neurosci 32:16880–16891 Available at: http://www.jneurosci.org/cgi/doi/10.1523/JNEUROSCI.3807-12.2012.

St. Onge JR, Floresco SB (2009) Dopaminergic modulation of risk-based decision making. Neuropsychopharmacology 34:681–697.

St. Onge JR, Floresco SB (2010) Prefrontal cortical contribution to risk-based decision making. Cereb Cortex 20:1816–1828.

Stopper CM, Green EB, Floresco SB (2014) Selective involvement by the medial orbitofrontal cortex in biasing risky, but not impulsive, choice. Cereb Cortex 24:154–162.

Sugam JA, Day JJ, Wightman RM, Carelli RM (2012) Phasic nucleus accumbens dopamine encodes risk-based decision-making behavior. Biol Psychiatry 71:199–205 Available at: http://dx.doi.org/10.1016/j.biopsych.2011.09.029.

van Gaalen MM, van Koten R, Schoffelmeer ANM, Vanderschuren LJMJ (2006) Critical Involvement of Dopaminergic Neurotransmission in Impulsive Decision Making. Biol Psychiatry.

Verdejo-Garcia A, Chong TT, Stout JC, Yucel M, London ED (2018) Stages of dysfunctional decision-making in addiction. Pharmacol Biochem Behav.

Wade TR, De Wit H, Richards JB (2000) Effects of dopaminergic drugs on delayed reward as a measure of impulsive behavior in rats. Psychopharmacology (Berl).

Winstanley CA, Theobald DEH, Dalley JW, Robbins TW (2005) Interactions between serotonin and dopamine in the control of impulsive choice in rats: Therapeutic implications for impulse control disorders. Neuropsychopharmacology 30:669–682 Available at: http://dx.doi.org/10.1038/sj.npp.1300610.

Zalocusky KA, Ramakrishnan C, Lerner TN, Davidson TJ, Knutson B, Deisseroth K (2016) Nucleus accumbens D2R cells signal prior outcomes and control risky decision-making. Nature 531:642–646 Available at: http://www.nature.com/doifinder/10.1038/nature17400.

